# A Type VI Secretion System in *Burkholderia* Species *cenocepacia* and *orbicola* Triggers Distinct Macrophage Death Pathways Independent of the Pyrin Inflammasome

**DOI:** 10.1101/2023.09.28.559184

**Authors:** Nicole A. Loeven, Arianna D. Reuven, Abigail P. McGee, Clarrisa Dabi, Bethany W. Mwaura, James B. Bliska

## Abstract

The *Burkholderia cepacia* complex contains opportunistic pathogens that cause chronic infections and inflammation in lungs of people with cystic fibrosis. Two closely related species within this complex are *Burkholderia cenocepacia* and the recently classified *Burkholderia orbicola. B. cenocepacia* and *B. orbicola* encode a type VI secretion system and the effector TecA, which is detected by the pyrin/caspase-1 inflammasome, and triggers macrophage inflammatory death. In our earlier study the pyrin inflammasome was dispensable for lung inflammation in mice infected with *B. orbicola* AU1054, indicating this species activates an alternative pathway of macrophage inflammatory death. Notably, *B. cenocepacia* J2315 and K56-2 can damage macrophage phagosomes and K56-2 triggers activation of the caspase-11 inflammasome, which detects cytosolic LPS. Here we investigated inflammatory cell death in pyrin-deficient (*Mefv*^−/−^) mouse macrophages infected with *B. cenocepacia* J2315 or K56-2 or *B. orbicola* AU1054 or PC184. Macrophage inflammatory death was measured by cleavage of gasdermin D protein, release of cytokines IL-1α and IL-1β and plasma membrane rupture. Findings suggest that J2315 and K56-2 are detected by the caspase-11 inflammasome in *Mefv*^−/−^ macrophages, resulting in IL-1β release. In contrast, inflammasome activation is not detected in *Mefv*^−/−^ macrophages infected with AU1054 or PC184. Instead, AU1054 triggers an alternative macrophage inflammatory death pathway that requires TecA and results in plasma membrane rupture and IL-1α release. Amino acid variation between TecA isoforms in *B. cenocepacia* and *B. orbicola* may explain how the latter species triggers a non-inflammasome macrophage death pathway.

## Introduction

The *Burkholderia cepacia* complex (Bcc) comprise a group of Gram-negative bacterial species that exist in the environment (1). The Bcc are opportunistic pathogens that cause infections in immunocompromised individuals including people with cystic fibrosis (pwCF) (1). The Bcc typically cause chronic infections associated with inflammation and exacerbations in adult pwCF. Chronic infections are difficult to eliminate due to intrinsic antibiotic resistance in the Bcc. Chronic Bcc infections can progress to an acute disease called cepacia syndrome that is characterized by necrotizing pneumonia, bacteremia, and sepsis. *B. cenocepacia* accounts for ∼45% of Bcc isolates from pwCF in the United States and is considered one of the most virulent in this group because of its strong association with cepacia syndrome (2, 3). In addition, *B. cenocepacia* infection is a contraindication for lung transplantation (4). *B. cenocepacia* subgrouping based on *recA* lineages has resulted in classification of genomovars IIIA-D (5) and shown that isolates belonging to genomovar IIIA commonly circulate in the UK, Europe, and Canada, while genomovar IIIB isolates more commonly cause infections in pwCF in the United States (6).

*B. cenocepacia* encodes numerous virulence factors (1, 7) and preferentially resides within phagocytes in the CF lung (8-10). Residence of *B. cenocepacia* in phagocytes has been observed by in situ imaging of human lung sections from transplant or autopsy, or mouse lung sections from experimental infections (11, 12). *B. cenocepacia* resides in vacuoles in phagocytic cells (11, 12), but may damage phagosomes and escape into the cytosol as well (13, 14). Additionally, *B. cenocepacia* can invade and survive in airway epithelial cells which may play a role in transmigration across the epithelial barrier (7). The intracellular location of *B. cenocepacia* appears to be key to its pathogenesis (8). *B. cenocepacia* can prevent phagosomal maturation and delay co-localization of the NADPH oxidase complex with the vacuolar membrane (15, 16). CF macrophages display defects in oxidative burst (17) and bacteria-specific autophagy (xenophagy) (9, 10) which may render them permissive to *B. cenocepacia.* In non-CF macrophages, phagosome damage by *B. cenocepacia* increases xenophagy (18) but the pathogen may be able to subvert this bactericidal process for intracellular survival (13). In addition, type I interferon production by non-CF macrophages in response to cytosolic *B. cenocepacia* infection can act in an autocrine fashion to increase xenophagy and limit bacterial replication (14).

A *B. cenocepacia* virulence factor implicated in lung infection and intracellular pathogenesis a type VI secretion system (T6SS) (8, 19). The T6SS contains a tube-like structure that functions to deliver effectors into bacteria or host cells (20, 21). Upon contact with target cells the tube comprised of the Hcp protein fires to puncture membranes and release associated effectors. The effectors have enzymatic activity that modify and inactivate key cellular processes or structures in target cells. The *B. cenocepacia* T6SS belongs to the first of three classes of these systems to be characterized (T6SS^i^, T6SS^ii^ and T6SS^iii^) (21). The T6SS^i^ class is common within Gammaproteobacteria and since the *B. cenocepacia* system is homologous to T6SS-1 of *Burkholderia pseudomallei* (22) we hereafter referred to it as T6SS-1. T6SS-1 is required for intracellular *B. cenocepacia* to inactivate small GTPases including Rac1 and Cdc42, resulting in disruption of the actin cytoskeleton and preventing assembly of the NADPH oxidase complex in macrophages (23, 24).

T6SS-1 is detected by inflammasomes in macrophages harboring *B. cenocepacia* (18, 25-27). Canonical caspase-1 inflammasomes are assembled when sensors such as NLRP3, NLRC4 or pyrin detect danger signals generated by bacterial pathogens during infection of host cells (28). Active caspase-1 is produced and cleaves the pro-inflammatory cytokine pro-IL-1β and the gasdermin-D (GSDMD) protein (29). Cleavage of GSDMD releases a domain that oligomerizes and forms pores in the plasma membrane (29). Mature IL-1β leaks out of these pores and can stimulate inflammatory signaling through activation of the IL-1 receptor found on a variety of host cells (29). Pore formation in plasma membranes can also result in pyroptosis, a pro-inflammatory form of cell death (29).

Early studies implicated T6SS-1 of *B. cenocepacia* strain K56-2 (genomovar IIIA, Table 1) in activation of the NLRP3 and NLRC4 inflammasomes in murine macrophages and the pyrin inflammasome in human macrophages (25, 26). Assembly of the pyrin inflammasome in response to T6SS-1 was subsequently observed in murine macrophages infected with J2315 (genomovar IIIA, Table 1) (27). Pyrin inflammasome assembly in macrophages infected with J2315 required inactivation of RhoA by T6SS-1 (27). The T6SS-1 in K56-2 has also been shown to be detected by the non-canonical caspase-11 inflammasome in murine macrophages (18, 30), which is consistent with phagosome damage (26) and entry of LPS into the cytosol (31). Caspase-11 can cleave GSDMD to form pores in macrophages but requires activation of a caspase-1 inflammasome (e.g. NLRP3) for processing and release of IL-1β (31).

**Table 1.**
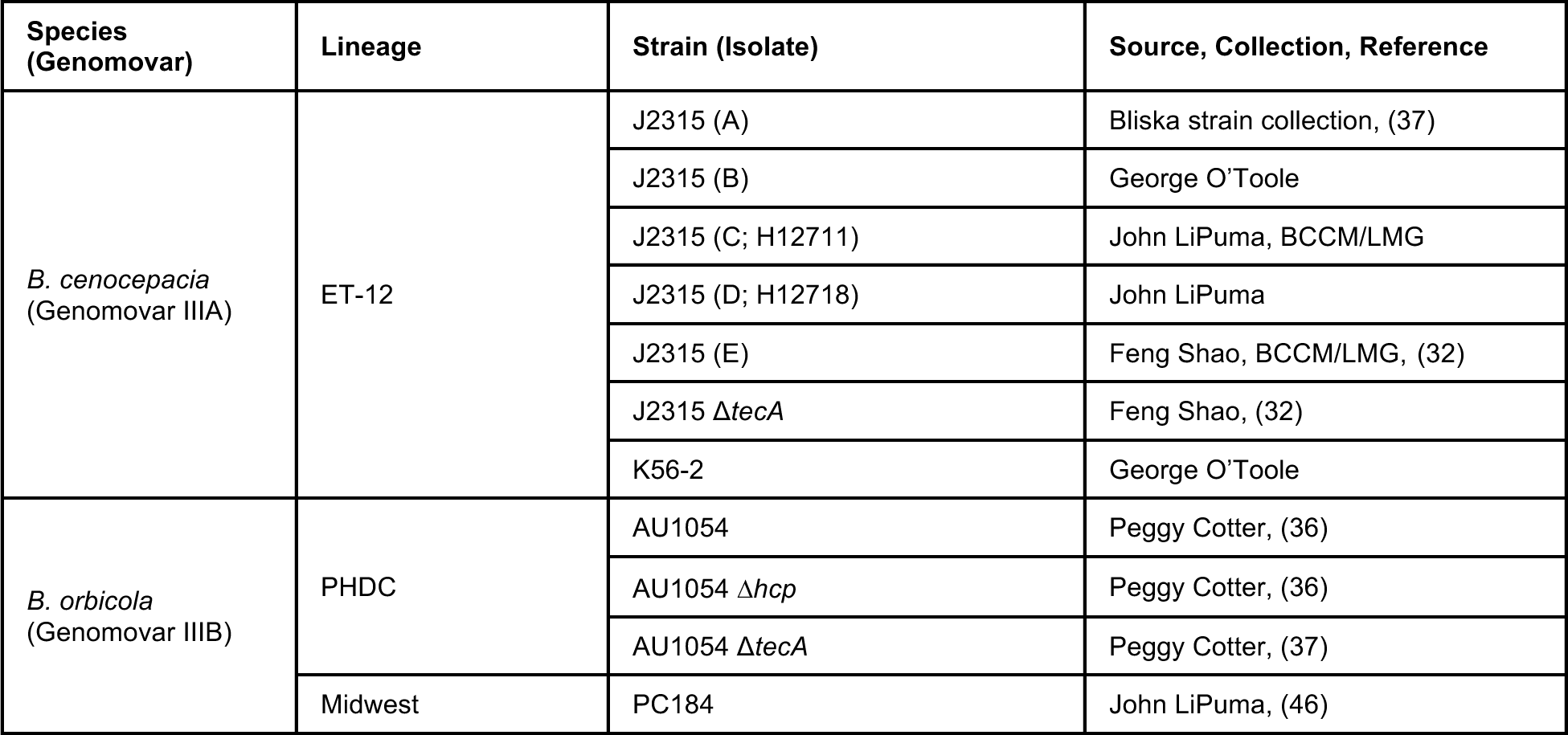
*B. cenocepacia* and *B. orbicola* isolates used in this study. Letters in parentheses after J2315 are arbitrary identifiers for isolates used in this study. H12711 and H12718 are additional identifiers for these isolates obtained from John LiPuma. Belgian Coordinated Collections of Microorganisms (BCCM/LMG)

Aubert et al. identified TecA (<u>T</u>6SS <u>e</u>ffector protein affecting <u>c</u>ytoskeletal <u>a</u>rchitecture) as being responsible for inactivation of RhoA and assembly of the pyrin inflammasome in macrophages infected with K56-2 or J2315 (32). In silico modeling of *B. cenocepacia* TecA predicted a fold consistent with a cysteine protease-like hydrolase, and putative catalytic C41, H105 and D149 residues were confirmed by mutagenesis (32). TecA appears to be the founding member of a T6SS effector superfamily with deamidase activity in bacterial opportunistic pathogens and symbionts (32). After entry into phagocytes *B. cenocepacia* employs the T6SS-1 to deliver TecA across the phagosome into the cytosol (32). RhoA is inactivated in response to deamidation of Asn-41 by TecA (32). Assembly of the pyrin inflammasome in response to deamidation of RhoA by TecA in macrophages infected with *B. cenocepacia* is an example of a guard mechanism of effector-triggered immunity (32). More recently TecA has been shown to promote localized actin polymerization that delays maturation of phagosomes containing J2315 in macrophages (33).

Previous studies examined the role of pyrin and TecA in lung pathogenesis by intranasally infecting C57BL/6 mice with J2315 (27, 32). Xu et al. compared C57BL/6 and pyrin knockout (*Mefv*^−/−^) mice and found that J2315 infection stimulated inflammatory cell infiltration into lungs of C57BL/6 but not *Mefv*^−/−^ mice (27). Aubert et al. infected C57BL/6 mice with J2315 or a Δ*tecA* mutant and reported that TecA was important for inflammatory cell infiltration and acute lung damage (32).

Increasing numbers of whole genome sequences for *Burkholderia* strains are being determined and analyzed. Results of two recent genomic analysis studies have suggested that genomovar IIIB isolates be designated as a species separate from *B. cenocepacia* (34, 35). Morales-Ruíz et al. propose for the new species the name *B. orbicola* (35). This name comes from the Latin word *orbis* and suffix-*cola* meaning “the whole world” and “dweller” respectively, making *orbicola* “inhabitant of the whole world” (35). Here, we follow this convention and hereafter refer to genomovar IIIB strains as *B. orbicola*.

*B. orbicola* strain AU1054 (Table 1), isolated from the bloodstream of a CF patient, encodes T6SS-1 (36) and TecA (37). Results of murine macrophage infections in vitro showed that AU1054 TecA triggers release of IL-1β in a T6SS-1- and pyrin-dependent manner (37). In an infection model using C57BL/6 or CF mice (mice with the most common CFTR mutation in humans, F508del) with AU1054 we confirmed that TecA is important for inflammatory cell recruitment to lungs (37). Surprisingly, using *Mefv*^−/−^ mice in the CF or non-CF background we were unable to reproduce a role for pyrin in inflammatory cell accumulation in lungs during infection with AU1054 or J2315 (37). This finding led us to hypothesize that AU1054 and J2315 can trigger inflammation by a mechanism independent of the pyrin inflammasome. Notably, J2315 been shown to damage macrophage phagosomes (13), which would allow detection by the non-canonical inflammasome, similar to its clonal relative K56-2 (18, 30). Activation of the caspase-11 inflammasome has been shown to contribute to lung inflammation and immunopathology in mice infected with K56-2 (18).

Here we investigated pyrin-independent macrophage inflammatory cell death in response to infection with *B. cenocepacia* J2315 or *B. orbicola* AU1054. Results indicate that J2315 and AU1054 trigger distinct pyrin-independent macrophage inflammatory death pathways with different inflammasome and TecA requirements. Amino acid differences between TecA protein sequences of *B. cenocepacia* and *B. orbicola* strains were identified, suggesting distinct effector isoform activities. These data have important implications for understanding the role of T6SS-1 and TecA in macrophage inflammatory death and the pathogenesis of chronic lung infections caused by *B. cenocepacia* and *B. orbicola* in pwCF.

## Results

### Pyrin is differentially required for gasdermin-D cleavage in macrophages infected with *B. cenocepacia* or *B. orbicola*

To determine if AU1054 and J2315 can trigger inflammasome activation independent of pyrin, bone marrow derived macrophages (BMDMs) from *Mefv^+/+^* or *Mefv*^−/−^ C57BL/6 mice were LPS primed, infected for 90 min and cleavage of GSDMD was measured by immunoblotting. For control strains we used AU1054 11*tecA* and 11*hcp* mutants which are unable to trigger the pyrin inflammasome (37) and K56-2 which activates the caspase-11 inflammasome (18, 30) (Table 1). As expected GSDMD cleavage in BMDMs infected with AU1054 required TecA, Hcp and pyrin (Fig.1, lanes 2-4, 8-10). Interestingly, GSDMD was cleaved in *Mefv^+/+^* and *Mefv*^−/−^ BMDMs infected with J2315 (isolate A, Table 1), like K56-2 (Fig.1, lanes 5,6 and 11,12). These results suggest that J2315 can, like K56-2, activate a pyrin-independent inflammasome in macrophages.

**Fig.1.**
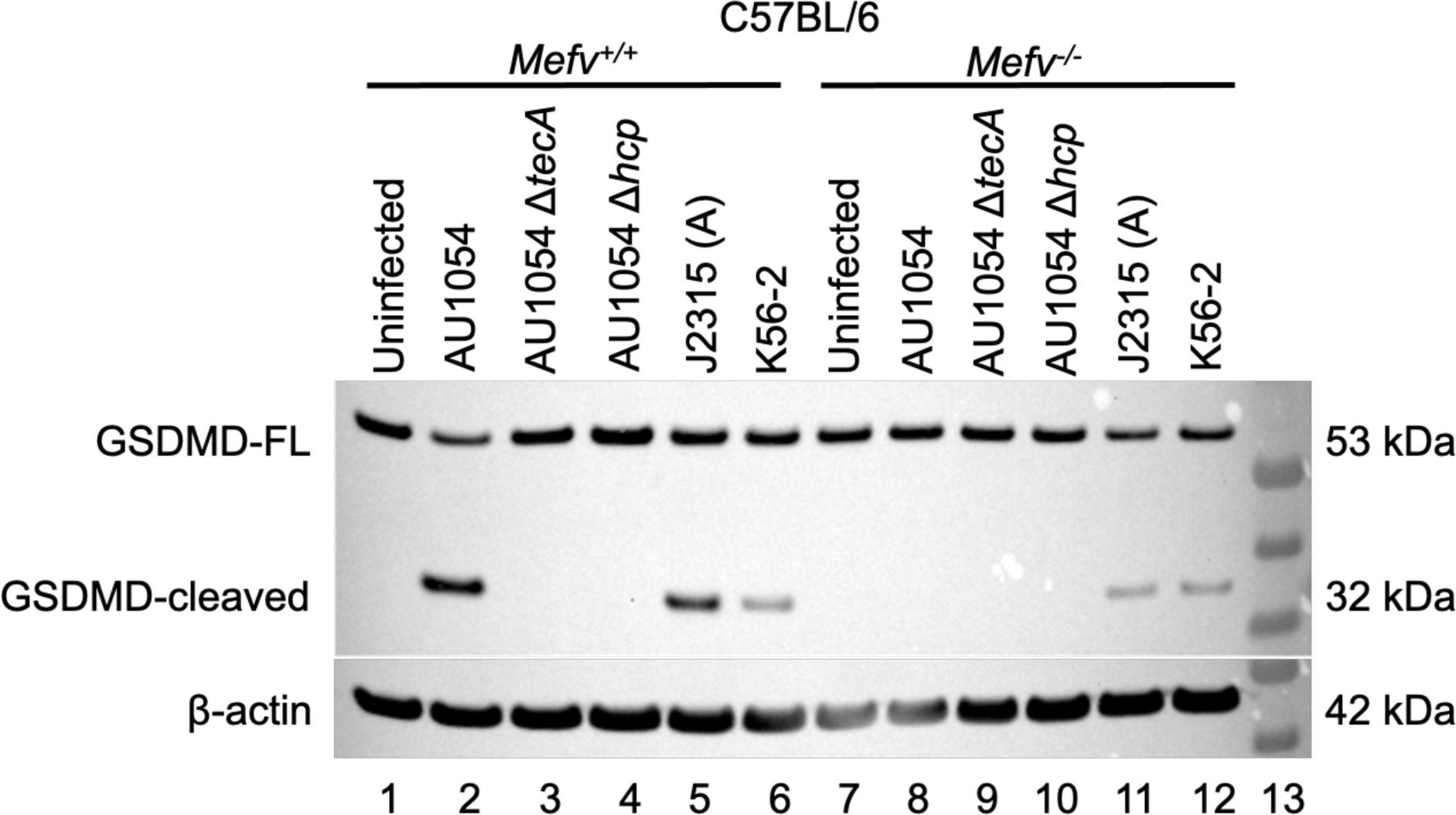
Pyrin is differentially required for GSDMD cleavage in BMDMs infected with AU1054, J2315 or K56-2. LPS-primed C57BL/6 *Mefv^+/+^* or *Mefv^−/−^* BMDMs were left uninfected or infected with the indicated strains (Table 1) for 90 min at MOI 20 and lysates were analyzed by immunoblotting with antibodies to GSDMD or β-actin. Positions of GSDMD 53 kDa full length (FL) and 32 kDa cleaved forms are indicated. Lane 13 is molecular weight (MW) ladder.

To determine if the phenotype we observed was specific to the J2315 A isolate from our laboratory, several additional isolates obtained from other investigators (Isolates B-D, Table 1) were analyzed in infections with *Mefv^+/+^*and *Mefv*^−/−^ BMDMs. GSDMD was cleaved in *Mefv^−/−^* BMDMs infected with all isolates (Fig.2A, lanes 8,9 and Fig.2B, lanes 7-9), indicating that pyrin-independent inflammasome activation and GSDMD cleavage is a general property of J2315.

**Fig.2.**
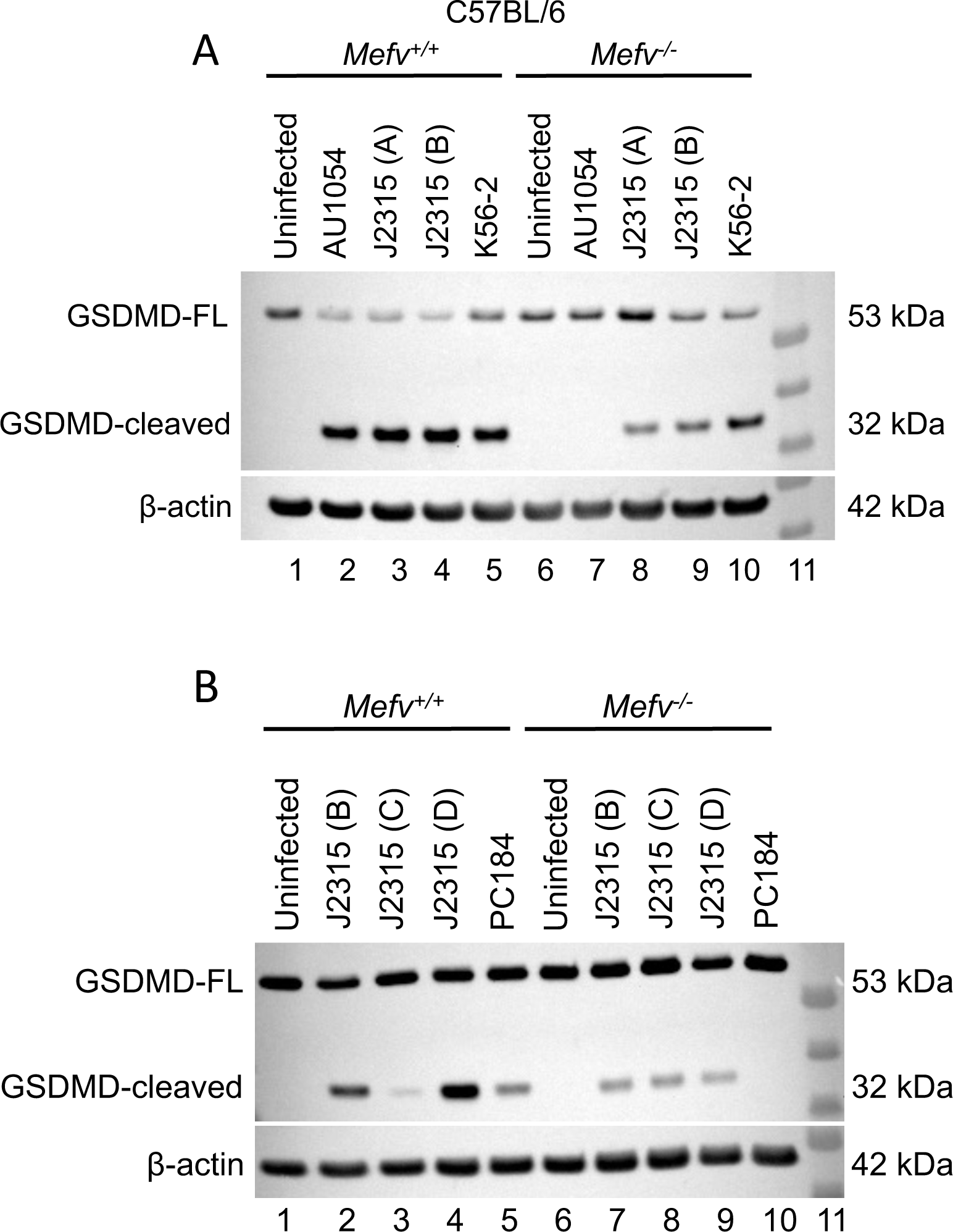
Pyrin is differentially required for GSDMD cleavage in BMDMs infected with *B. cenocepacia* and *B. orbicola*. LPS-primed C57BL/6 *Mefv^+/+^*and *Mefv^−/−^* BMDMs were left uninfected or infected for 90 min at MOI 20 with the indicated strains (Table 1) and lysates were analyzed by immunoblotting with antibodies to GSDMD or β-actin. The result in (B) with PC184 is representative of two independent experiments. Lanes 11 are a MW ladder.

We also infected *Mefv^+/+^* and *Mefv*^−/−^ BMDMs with an additional *B. orbicola* isolate, PC184 (Table 1), and found that pyrin was required for cleavage of GSDMD (Fig.2B, lanes 5,10), like AU1054. These results suggest that *B. orbicola* and *B. cenocepacia* differ in the ability to activate inflammasomes in macrophages independent of pyrin.

### TecA is dispensable for pyrin-independent GSDMD cleavage and IL-1β release in macrophages infected with J2315

To determine if TecA is required for J2315 to trigger pyrin-independent GSDMD cleavage and IL-1β release, we obtained an additional isolate (Isolate E) of this strain and its corresponding Δ*tecA* mutant (32) (Table 1). LPS-primed *Mefv^+/+^* or *Mefv^−/−^*BMDMs infected with these strains or AU1054 controls were analyzed for GSDMD cleavage as well as IL-1β release. As expected from our previous ELISA results (37) and the GSDMD cleavage data presented here (Figs.1, 3A), TecA and pyrin were required for significant release of IL-1β from BMDMs infected with AU1054 (Fig.3B). However, findings here showed that TecA and pyrin were not required for GSDMD cleavage (Fig.3A, lane 11) or IL-1β release BMDMs infected with J2315 (Fig.3C). Additional experiments are needed to solidify these findings because the IL-1β values for the J2315 samples showed variability. Nevertheless, these results are consistent with the idea that J2315, like K56-2, can activate both pyrin and the caspase-11 inflammasomes in macrophages (18). This conclusion will need to be verified using *Casp11^−/−^* BMDMs in future experiments.

**Fig.3.**
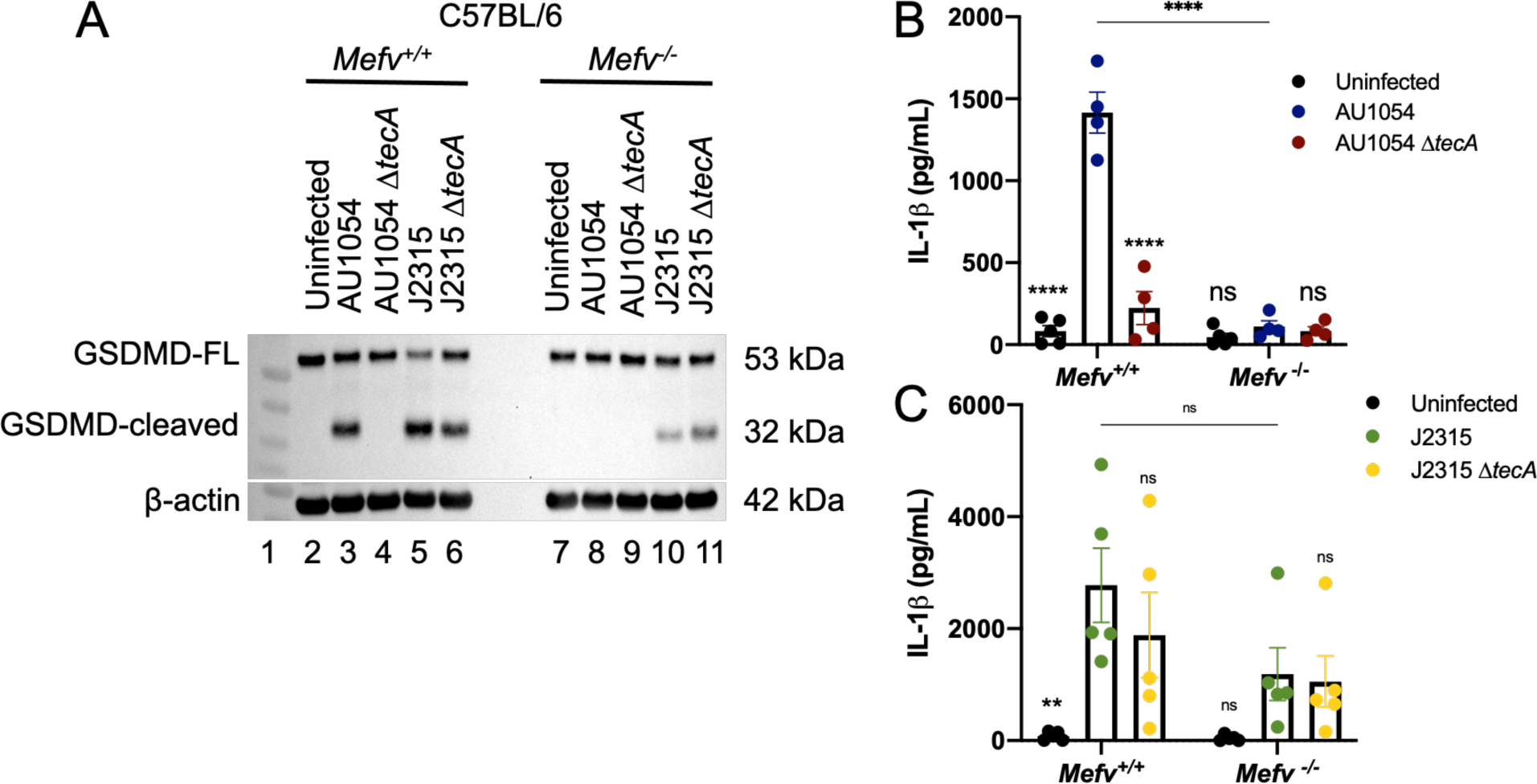
TecA is dispensable for pyrin-independent GSDMD cleavage and IL-1β release in BMDMs infected with J2315. LPS-primed C57BL/6 *Mefv^+/+^* or *Mefv^−/−^* BMDMs were left uninfected or infected with the indicated strains (Table 1) at an MOI of 20 for 90 minutes. (A) BMDM lysates were analyzed by immunoblotting with antibodies to GSDMD or β-actin. Lane 1 is a MW ladder. Cell supernatants were collected and analyzed for IL-1β by ELISA (B,C). Data shown in (B,C) are from four or five independent experiments, respectively. Error bars represent standard deviation. Significance determined by two-way ANOVA and Tukey posttest as compared to AU1054 (B) or J2315 (C) within and between BMDM genotypes. ns, not significant;

### TecA triggers pyrin inflammasome-independent IL-1α and LDH release from macrophages infected with AU1054

To determine if AU1054 triggers another form of pyrin-independent macrophage inflammatory death, we measured release of IL-1α and LDH. IL-1α can be released from macrophages by inflammasome-dependent and -independent mechanisms. In the latter case plasma-membrane rupture (PMR) allows IL-1α to be released from host cells to act as a danger-associated-molecular pattern (DAMP) (38). LDH at 140 kDa is a marker of PMR. These experiments used *Mefv^+/+^* or *Mefv^−/−^* BMDMs from mice engineered with a CF genotype (*Cftr^F508del^*) and after LPS priming they were left uninfected or infected with AU1054 or AU1054 11*tecA* as above. For a positive control we treated the BMDMs with *Clostridium difficile* TcdB toxin and the culture supernatants were analyzed by ELISA for IL-1β, which is dependent upon inflammasome activation for processing and release. As shown in Fig.4A, significant IL-1β release upon AU1054 infection required TecA and pyrin as expected. However, pyrin was not required for TecA to trigger release of IL-1α (Fig.4B) or LDH (Fig.4C) from *Mefv^−/−^* BMDMs infected with AU1054. IL-1α released in response to AU1054 infection was significantly lower in *Mefv^−/−^* as compared to *Mefv^+/+^* or BMDMs (Fig.4B), likely because pyrin inflammasome activation contributed to secretion of the cytokine in the former case. IL-1α and LDH released from *Mefv^−/−^* BMDMs intoxicated with TcdB was at background levels (Fig.4B, C respectively). These data suggest that AU1054 TecA triggers an inflammasome-independent mechanism of PMR leading to macrophage inflammatory death.

**Fig.4.**
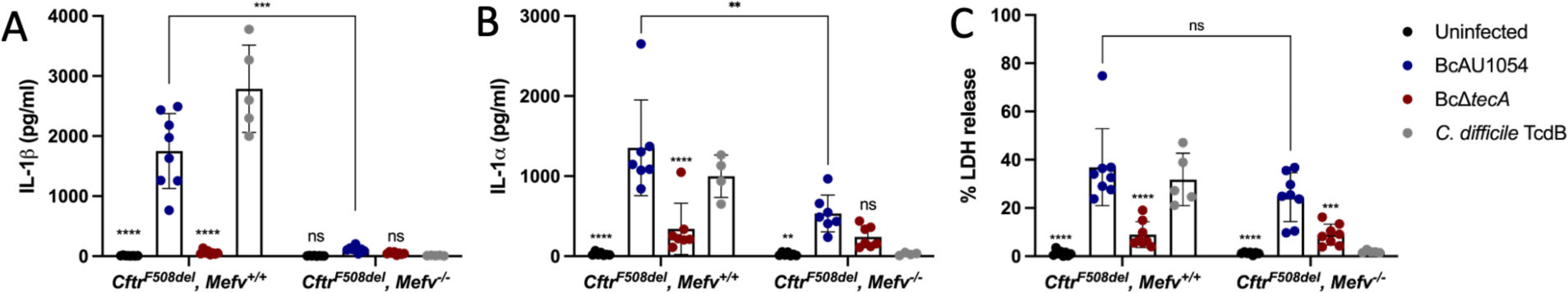
TecA triggers pyrin-independent IL-1α and LDH release from BMDMs infected with AU1054. LPS-primed *Cftr^F508del^ Mefv^+/+^* or *Mefv^−/−^*BMDMs were left uninfected or infected with AU1054 or AU1054 Δ*tecA* at an MOI of 20 or intoxicated with TcdB for 90 minutes. Cell supernatants were collected and analyzed for IL-1β (A) or IL-1α (B) by ELISA and PMR by LDH release (C). Data shown are from at least three independent experiments. Error bars represent standard deviation. Significance determined by two-way ANOVA as compared to AU1054 within and between BMDM genotypes. ns, not significant; ** p <0.01, *** p <0.001, **** p <0.0001.

### The TecA proteins of *B. orbicola* and *B. cenocepacia* exhibit amino acid variation

The ability of AU1054 TecA to trigger inflammasome-independent PMR in BMDMs (Fig.4) suggested that the *orbicola* isoform has distinct biological function/activity as compared to the canonical *cenocepacia* effector. To examine this possibility, we sequenced *tecA* genes from isolates used in this study. The resulting predicted TecA protein sequences were aligned using ClustalW (Fig.5). The *B. cenocepacia* J2315 and K56-2 proteins are 100% identical and the *B. orbicola* AU1054 and PC184 proteins are 99% identical (Fig.5). In contrast, *B. cenocepacia* TecA shares only ∼90% sequence identity with the *B. orbicola* isoform, due to 15-16 amino acid differences (Fig.5).

**Fig.5.**
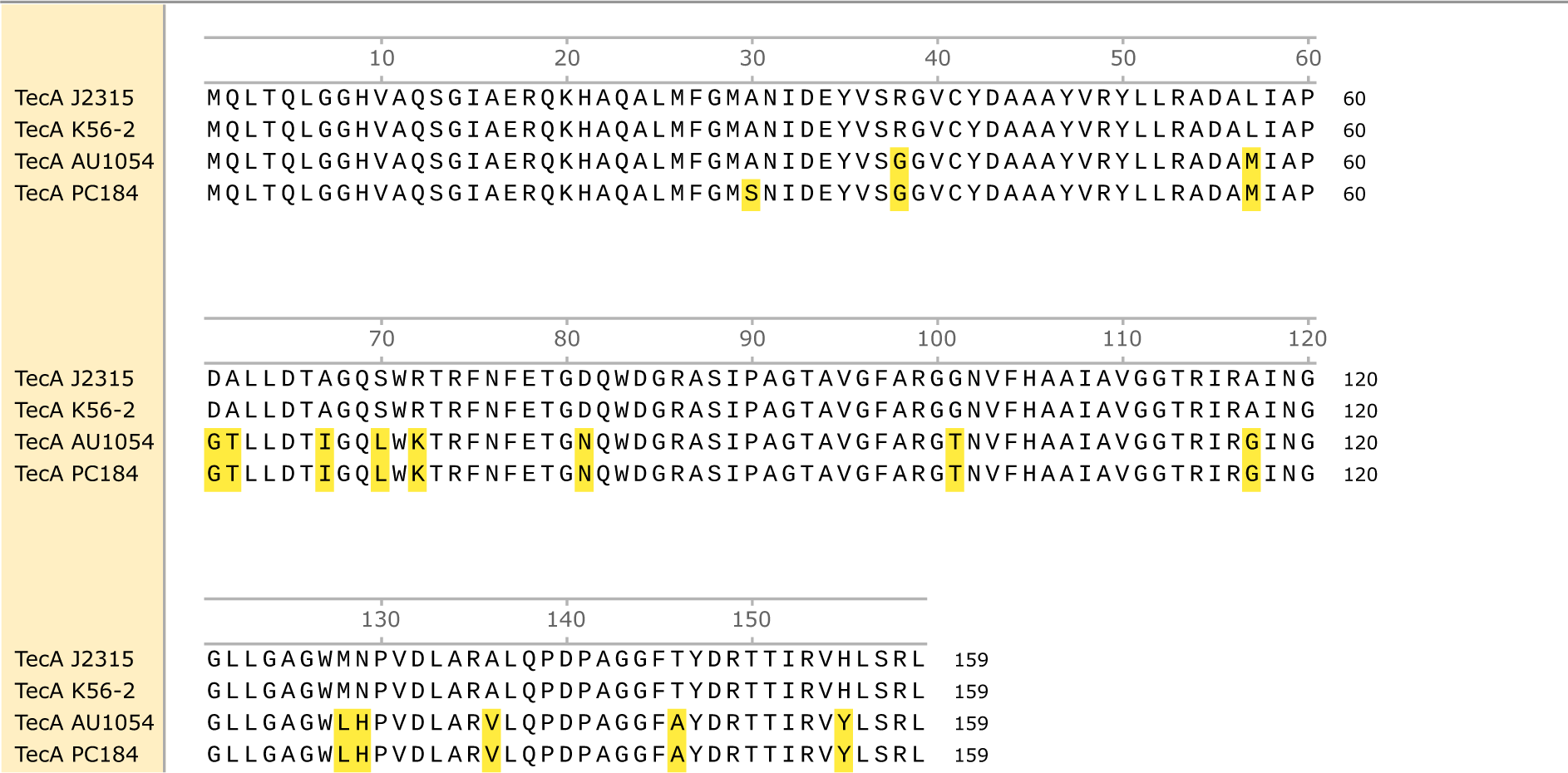
The TecA proteins of *B. cenocepacia* and *B. orbicola* exhibit amino acid variation. The *tecA* genes of the strains used in this study were sequenced and then translated and aligned in SnapGene using ClustalW. Amino acids highlighted in yellow indicate differences from the TecA sequence of J2315.

All TecA isoforms have residues C41, H105 and D148 that are important for catalytic activity (32, 37) (Fig.5). In addition, amino acids immediately flanking the catalytic residues are highly conserved between *B. cenocepacia* and *B. orbicola* (Fig.5). However, *B. orbicola* TecA has non-conserved substitutions R38G (charged to non-charged), G101T (non-polar to polar), and T146A (polar to non-polar) within 3, 4 and 2 amino acids of C41, H105 and D148, respectively (Fig.5).

Predicted structures of the K56-2 and AU1054 TecA isoforms were determined by AlphaFold2 and displayed by PyMol (Fig.6). Results show that the overall folds and positions of catalytic residues are highly similar (Fig.6). However, the presence of the non-conserved substitutions R38G (charged to non-charged), G101T (non-polar to polar), and T146A (polar to non-polar) in the vicinity of the active site (Fig.6) could impact the biological activity of the *B. orbicola* TecA protein.

**Fig.6.**
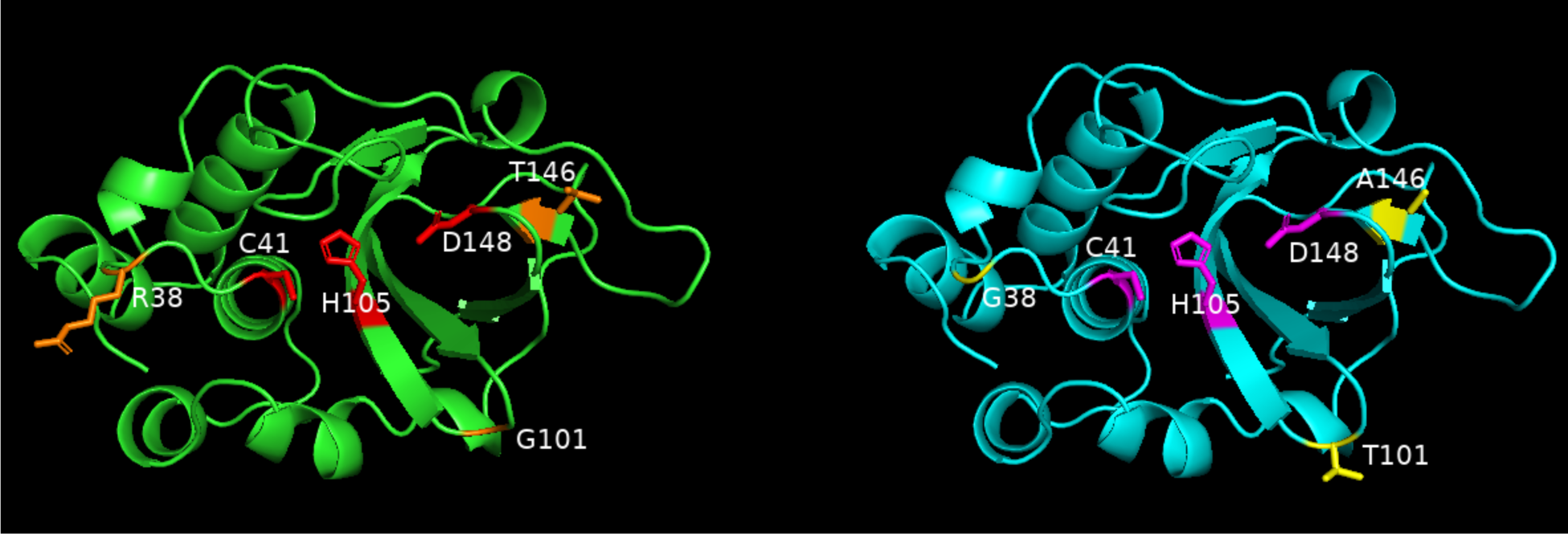
Comparison of predicted structures of TecA proteins of *B. cenocepacia* and *B. orbicola* and non-conserved amino acids adjacent to catalytic sites. Predicted structures of TecA proteins of K56-2 (left) and AU1054 (right) were generated using AlphaFold2 and displayed using PyMol. Catalytic residues are shown in red (left) or cyan (right). Non-conserved residues adjacent to catalytic sides are shown in orange (left) or yellow (right).

## Discussion

The data presented above increase our understanding of the role of T6SS-1 and TecA in the pathogenesis of *B. orbicola* and *B. cenocepacia* in the following three ways. First, our results confirm that *B. cenocepacia* triggers inflammasome-dependent macrophage inflammatory death by two mechanisms, one that is TecA and pyrin dependent and the other likely requires detection of vacuole damage by caspase-11 (18). Second, *B. orbicola* via T6SS-1 triggers TecA-dependent macrophage inflammatory death by two mechanisms, one that requires pyrin and the other is inflammasome independent. Third, the TecA protein of *B. orbicola* appears to comprise a separate subgroup of this effector superfamily, with distinct biological activity. Together, these results indicate that the underlying mechanism of inflammation in chronic *B. orbicola* and *B. cenocepacia* lung infections is multifactorial and species specific, which has implications for treating this disease in pwCF.

J2315 and K56-2 are genomovar IIIA CF clinical isolates and members of the epidemic ET12 lineage common in Europe (39). Despite being clonal, the literature indicates that during macrophage infections the T6SS-1 of J2315 is only detected by the pyrin inflammasome via TecA (27, 32) while K56-2 is detected by the pyrin and caspase-11 inflammasomes (18, 25, 30). Our results address this discrepancy by showing that when analyzed in parallel J2315 and K56-2 both trigger pyrin-independent GSDMD cleavage in infected BMDMs. In addition, J2315 causes low but significant IL-1β release from *Mefv*^−/−^ BMDMs under our infection conditions. It is probable that caspase-11 is the proximal inflammasome activated in *Mefv*^−/−^ BMDMs infected with J2315 and K56-2. It will be important to carry out *B. cenocepacia* infection experiments with BMDMs deficient in caspase-11 to confirm this hypothesis. Activation of caspase-11 by J2315 may not have been detected previously (27, 32) due to insufficient lengths of LPS priming of *Mefv*^−/−^ BMDMs. The two hour LPS priming used by Xu et al. and Aubert et al. (27, 32) may have been insufficient to activate expression of caspase-11 (31). The overnight LPS priming we use to activate expression of pyrin may have fortuitously allowed us to detect activation of caspase-11 in *Mefv*^−/−^ BMDMs infected with J2315 and K56-2.

Xu et al. found that lung inflammation in mice infected with J2315 required pyrin (27). However, in preliminary experiments we observed only a subtle decrease in lung inflammation in mice infected with J2315 (37). Experimental variables such as genetic or microbiota differences between *Mefv*^−/−^ mouse lines used by Xu et al. (27) and our study (37) could be responsible for the discrepancy. However, our result can be explained by the data of Krause et al. (18) who showed that caspase-11 is important for lung inflammation in mice infected with K56-2 (18). Presumably, inflammation triggered by caspase-11 could partially compensate for the absence of pyrin in *Mefv*^−/−^ mice infected with J2315. It is possible that pyrin and caspase-11 contribute additively to inflammation in lungs during *B. cenocepacia,* and this could be studied by directly comparing responses to *B. cenocepacia* infection in mice that are *Mefv*^−/−^ or *Casp11^−/−^*or deficient in both inflammasomes. Interestingly, *Casp11^−/−^* mice had increased survival compared to *Casp1^+/+^* mice, even though K56-2 growth was restricted better in the latter animals (18). This result indicates that lethality in this context is due to immunopathology in the lung rather than bacterial growth in per se.

The mechanism of caspase-11 activation by J2315 in macrophages is likely the same as K56-2, where T6SS-1 has been shown to mediate phagosome damage (26). In fact, there is evidence that J2315 escapes from damaged phagosomes (13, 14). The mechanism of phagosome damage by *B. cenocepacia* T6SS-1 is unknown. Polymerized actin co-localizes with cytosolic J2315 in human macrophages (13). However, the role of actin polymerization in phagosome damage by *B. cenocepacia* is unclear. Additional evidence that *B. cenocepacia* escapes phagosomes comes from the finding that J2315 activates non-inflammasome PRRs like mitochondrial antiviral-signaling protein (MAVS) and stimulator of interferon genes (STING) in macrophages (14). MAVS is activated primarily in response to viral RNAs, where STING is activated in response to detection of bacterial double stranded DNA (dsDNA) (14). MAVS and STING activation in macrophages leads to a type I interferon (IFN) response which induces autophagy/xenophagy and was shown to counteract *B. cenocepacia* J2315 replication in macrophages (14).

Our data suggest that *B. orbicola* TecA can trigger PMR in infected macrophages by an inflammasome-independent mechanism because the lack of GSDMD cleavage in *Mefv*^−/−^ macrophages indicates that caspase-1 and caspase-11 are not being activated. *B. orbicola* TecA can trigger release of LDH from unprimed BMDMs (unpublished observation) further suggesting this pathway of PMR is inflammasome independent. To confirm this concept, it will be important to carry out *B. orbicola* infection experiments with BMDMs deficient in caspase-1 and −11. Release of HMGB1 could be measured to confirm that multiple DAMPs are being released by PMR. It will also be important to determine if *B. orbicola* TecA triggers inflammasome-independent PMR in human macrophages.

Recently the protein NINJ1 has been shown to be required for PMR downstream of multiple forms of cell death in BMDMs (40). It is possible that NINJ1 is important for inflammasome-independent PMR and release of DAMPs from *B. orbicola*-infected macrophages. Microscopy could be used to confirm that inflammasome-independent DAMP release is associated with plasma membrane damage in *Mefv*^−/−^ macrophages infected with *B. orbicola*. Propidium iodide uptake measured by fluorescence microscopy and cell rounding and blebbing as detected by phase microscopy would be indicators of plasma membrane damage (41).

The existence of an inflammasome-independent PMR pathway in macrophages likely explains why recruitment of inflammatory cells to lungs of mice infected with AU1054 is not decreased in the absence of pyrin (37). We hypothesize *B. orbicola* TecA triggers redundant pathways of macrophage inflammatory cell death both in vitro and in vivo. The pyrin-dependent pathway results in IL-1β release and pyroptosis, while the inflammasome-independent pathway results in PMR and release of multiple DAMPs. Either pathway can exacerbate inflammatory monocyte recruitment and persistence in lungs infected with *B. orbicola* (37). For example, IL-1β or DAMPs released from *B. orbicola*-infected alveolar macrophages could activate airway epithelial cells to produce chemokines for inflammatory monocytes. Inflammatory monocytes use the receptor CCR2 to migrate out of the bone marrow into sites of infection in response to the chemokine CCL2 (42). Inflammatory monocytes can differentiate into macrophages and depending on the context can cause detrimental immune responses in lung infections (42, 43). Persistence of excessive numbers inflammatory monocyte-derived macrophages could cause immunopathology in lungs infected with *B. orbicola* (37). Inflammatory monocyte-derived macrophages recruited by CCL2 could also become infected with *B. orbicola* and release more IL-1β or DAMPs, thus initiating a damaging cycle of inflammation.

Our finding that *B. orbicola* TecA triggers inflammasome-independent PMR suggests that the 10% amino acid variation in this isoform imparts different biological activity as compared to *B. cenocepacia*. Although speculative we suggest that *B. orbicola* TecA deaminates a host protein target which triggers PMR. To test the above hypothesis, it would be interesting to replace TecA in AU1054 with the *B. cenocepacia* isoform and determine how this allele swap impacts macrophage inflammatory death mechanisms upon bacterial infection.

## Acknowledgments

We thank George O’Toole, John LiPuma, Feng Shao and Sergio Grinstein for providing bacterial strains. Ericka A. Tamayo-Guevara helped produce the TecA AlphaFold structures shown in Fig.6. This work was supported by grants from the Cystic Fibrosis Foundation (BLISKA22G0 and STANTO19R0) and support by DartCF through NIH grant P30-DK117469.

## Materials and Methods

### Bacterial strains

Bacterial strains used in this study are shown in Table 1. *B. orbicola* and *B. cenocepacia* strains were grown at 37°C on Luria-Bertani (LB) agar plates or in LB both with shaking.

### Ethics statement

**I**solation of bone barrow cells from mice was carried out in accordance with a protocol that adhered to the Guide for the Care and Use of Laboratory Animals of the National Institutes of Health (NIH) and was reviewed and approved (approval #00002184) by the Institutional Animal Care and Use Committee at Dartmouth College. The Dartmouth College animal program is registered with the U.S. Department of Agriculture (USDA) through certificate number 12-R-0001, operates in accordance with Animal Welfare Assurance (NIH/PHS) under assurance number D16-00166 (A3259-01) and is accredited with the Association for Assessment and Accreditation of Laboratory Animal Care International (AAALAC, accreditation number 398).

### Mouse strains

Wild-type (WT) C57BL/6J (stock# 000664) mice were purchased from Jackson Laboratories. Pyrin-knockout mice (*Mefv^−/−^*) on the C57BL/6 background were obtained from Jae Chae and Daniel Kastner at the NIH (44) and bred in the mouse facilities at Dartmouth. Mice with the *Cftr^F508del^* mutation on the C57BL/6 background were obtained from Case Western Reserve University’s Cystic Fibrosis Mouse Models Core and bred at Dartmouth (37). *Mefv^−/−^* and *Cftr^F508del^*mice were crossed to obtain breeding pairs that were *Cftr^F508del^ Mefv^+/-^* and *Cftr^F508del^ Mefv^−/−^*. The resulting pups were genotyped as described (37) and used as a source of bone marrow cells.

### Preparation of BMDMs

Bone marrow derived macrophages (BMDMs) were cultured from bone marrow of mice and cultured as described previously (37). After 7 days of differentiation the BMDMs were seeded at a density of 0.8×10^6^ cells/well in 6-well plates in MGM 10/10 media containing Dulbecco’s Modified Eagle Medium (DMEM) + Glutamax (Gibco^®^) containing 10% Fetal Bovine Serum (FBS) (GE^®^), 10% L929 cell-conditioned media, 1 mM sodium pyruvate (Gibco^®^), 10 mM HEPES (Gibco^®^) and divided into 6-well plates at a density of 0.8 ×10^6^ cells/well in a total volume of 3 mL. The BMDMs were primed with 100 ng/mL O26:B6 *Escherichia coli* LPS (Sigma^®^) and incubated overnight at 37°C with 5% CO_2_.

### BMDM infection assays

Overnight (16-hour) cultures of *B. orbicola* or *B. cenocepacia* were subcultured 1:100 in fresh LB to OD_600_ = ∼0.05 on infection day and shaken at 37°C until the cultures reached mid-log phase (OD_600_ = 0.300). Cultures were then pelleted, the LB was removed, and the bacteria were resuspended in PBS to the original volume (∼3×10^8^ CFU/mL). The bacterial suspensions were then diluted to an MOI of 20 in warmed, in media lacking 10% FBS (MGM 0/10). As a positive control for pyrin inflammasome activation, the glucosyltransferase toxin from *Clostridium difficile* TcdB (List Biological Laboratories, Inc.) was also diluted to 0.1mg/ml in warmed MGM 10. The BMDMs were washed once in warm 1x PBS and 3ml fresh MGM 10 was added with bacteria or TcdB to the appropriate wells. The plates were then centrifuged for 5 minutes at 1,000rpm to bring the bacteria and the cells in contact on the bottom of the wells. The plates were incubated at 37°C with 5% CO_2_ for 90 minutes. Cell supernatants were collected for cytokine ELISAs and lactate dehydrogenase (LDH) assays. The BMDMs were lysed using mammalian protein extraction reagent (M-PER, Thermo Scientific^®^) with added cOmplete, Mini (Roche^®^) protease inhibitor and PhosSTOP (Roche^®^) phosphatase inhibitor (37).

### Western blotting

Five - 10mg of protein from the cell lysates were run on 4-12% NuPAGE Bis-Tris SDS-PAGE gels (Invitrogen by ThermoFisher Scientific^®^) and transferred to PVDF membranes (ThermoFisher Scientific^®^) using an iBlot 2 Gel Transfer Device (Life Technologies). Membranes were blocked in 5% non-fat dairy milk and incubated with primary antibody overnight. The primary antibodies used were rabbit MAb for GSDMD (Abcam^®^, ab209845), and rabbit polyclonal for b-actin (Cell Signaling^®^, #4967). HRP-conjugated anti-rabbit (Jackson laboratory) was used as a secondary antibody. Protein bands reacting with antibodies were visualized using chemiluminescent detection reagent (GE Healthcare^®^) on an iBright FL1500 (ThermoFisher Scientific^®^).

### IL-1β and IL-1α quantification

Murine IL-1β and IL-1α in BMDM supernatants was quantified using ELISA kits (R&D Systems^®^, MLB00C and MLA00, DLB50, respectively) following the manufacturer’s instructions.

### LDH quantification

LDH in BMDM supernatants was quantified using the CytoTox 96® Non-Radioactive Cytotoxicity Assay (Promega®) following the manufacturer’s instructions.

### Statistical analysis of data

Experimental data generated from at least three independent in vitro experiments are analyzed for significance. Probability (*p*) values for ELISA and LDH data are calculated using one-way analysis of variance (ANOVA) or grouped two-way ANOVA with Tukey’s posttest.

### TecA gene sequencing and protein sequence analysis

*B. orbicola* and *B. cenocepacia* were grown on LB agar plates, single colonies were picked, suspended in sterile Elution Buffer (Qiagen), and boiled for 5 minutes to release bacterial DNA. PCR was performed using the colony DNA and primers that anneal upstream (Forward 5’-ACGCCGTGACGGCCTGACGCCGT-3’) and downstream (Reverse 5’-GCGCGACGCCATGCGGAAATCGCCG-3’) of all four *tecA* genes analyzed (BCAM1857 in J2315, WQ49_RS04990 in K56-2, BCEN_RS18170 in AU1054 and EAY64595 in PC184). PCR products were run on a 1% agarose gel and purified using a Qiagen kit. Purified DNA was sequenced and resulting sequences were analyzed, translated, and aligned on SnapGene using ClustalW. Percent amino acid sequence identity was determined using NCBI Protein BLAST. Predicted structures of the AU1054 and K56-2 TecA isoforms were determined by AlphaFold2 (45) and displayed using PyMol.

